## Supplementary Figures 1-13 for "Evolutionary Origins and Functional Diversification of Auxin Response Factors"

#### Supplementary Figure 1

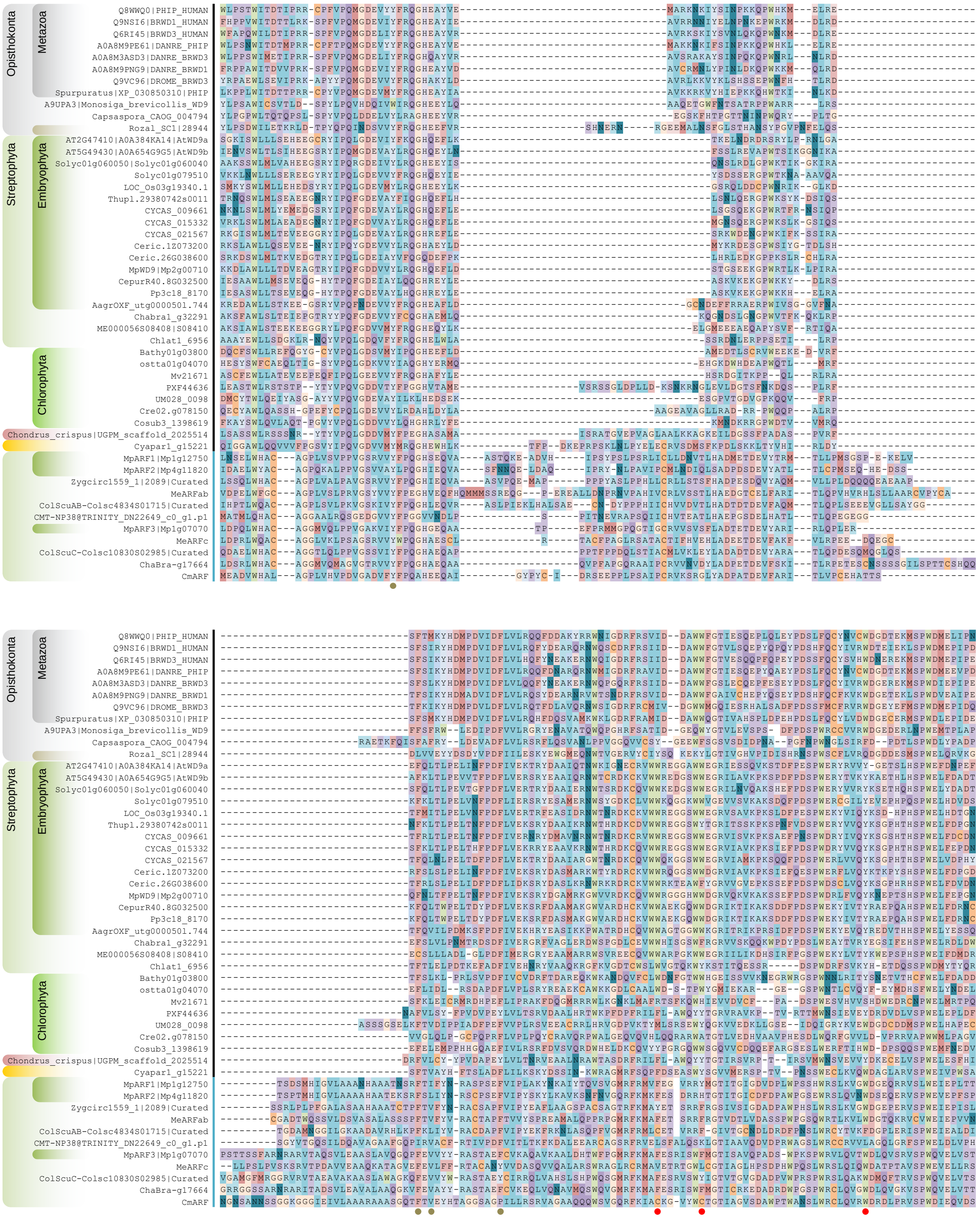


**Supplementary figure 1. PHIP-ARF homology at the primary sequence level**. Multiple sequence alignment of the cryptoTudor and DD-AD region of PHIP and ARF proteins. ARF B3 region has been cut from residues 119 to 250 (in MpARF1 as reference). Vertical black and cyan lines denote PHIP and ARF sequences, respectively. Filled circles denote predicted or putative hydrophobic cage residues in the DD or AD regions (tan brown and red, respectively). Note that despite HC conservation, DD cryptoTudor region in PHIP has not been found to hold histone binding activity as opposed to AD.

#### Supplementary Figure 2

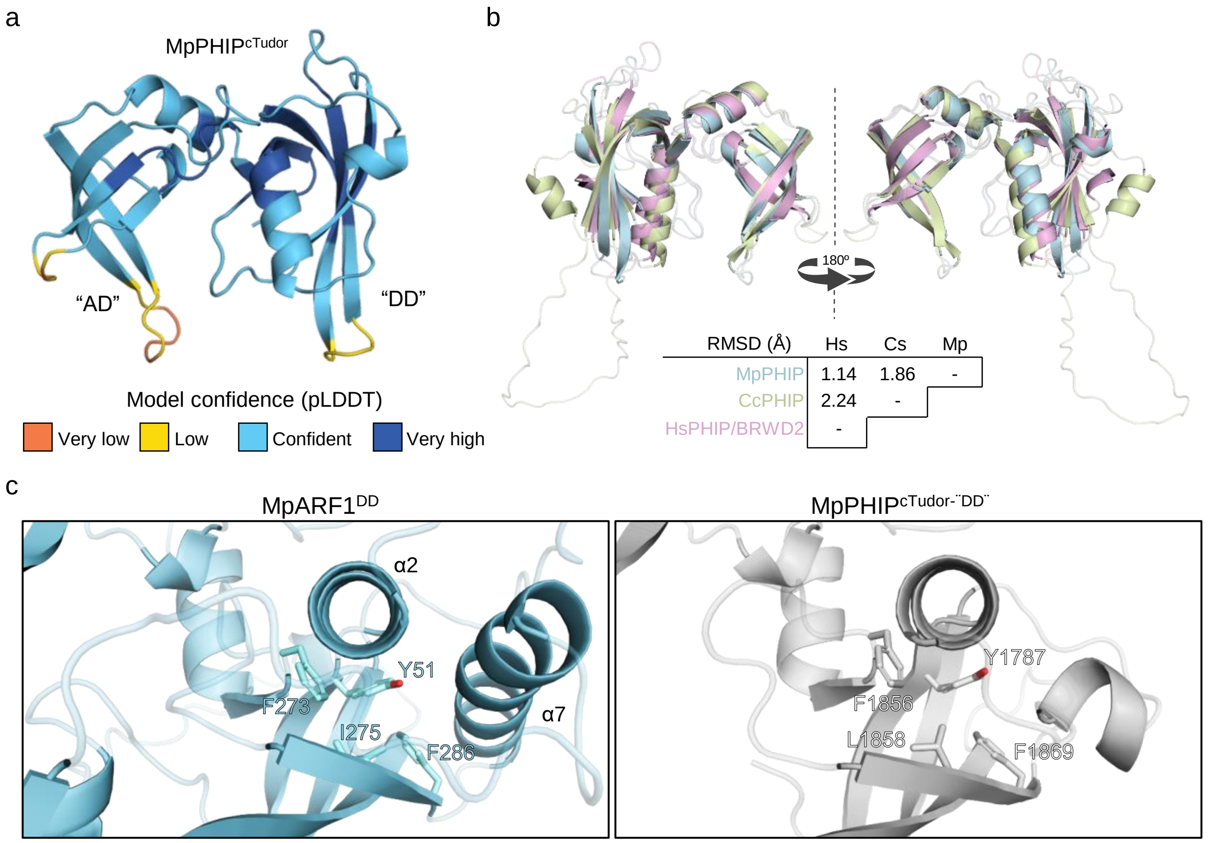


**Supplementary figure 2. PHIP crytoTudor and ARF DD-AD structural comparison**. **a**, AlphaFold2 structural prediction of MpPHIP cryptoTudor domain showing the confidence score per residue in the structure model. **b**, Structural alignment between different PHIP cTudors and root mean square deviation of pairwise alignments expressed in ångströms (Å) . MpPHIP, *Marchantia polymorpha*, blue; CcPHIP, *Chondrus crispus*, yellow; HsPHIP, *Homo sapiens*, pink. **c**, detail of the predicted hydrophobic cage in MpARF1 and MpPHIP dimerization domain (AD).

#### Supplementary Figure 3

a

| Construct | Lines | Postembryonic rescue (%) | |
|---|---|---|---|
| | | Yes (full) | No |
| *pARF5:ARF5* | 7 | 100 | 0 |
| *pARF5:ARF5$^{\Delta AD}$* | 7 | 0 | 100 |

b

| Construct | Lines | Growth rescue (%) | | Auxin response rescue (%) | |
|---|---|---|---|---|---|
| | | Yes | No | Yes | No |
| *pMpARF1:MpARF1* | 4 | 100 | 0 | 100 | 0 |
| *pMpARF1:MpARF1$^{\Delta AD}$* | 4 | 0 | 100 | 100 | 0 |

c

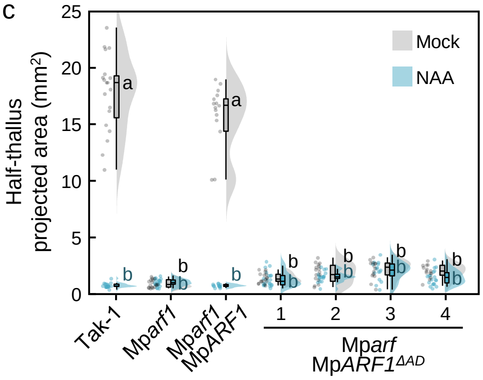


d

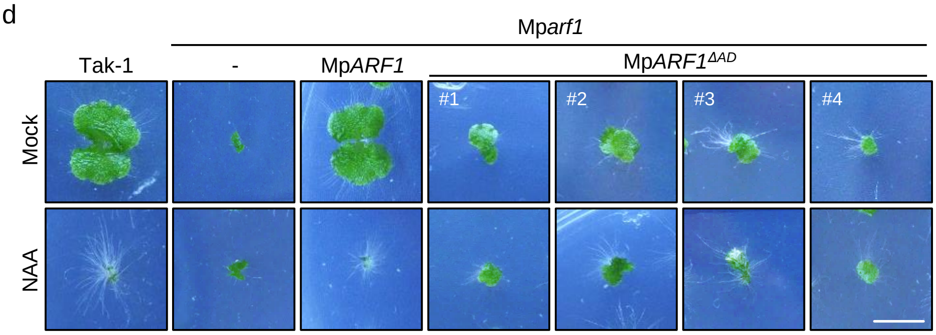


**Supplementary figure 3. AD region is essential for ARF function**. **a**, summary of Arabidopsis *mp* complementation assay using ARF5 with AD deleted (ARF5$^{AD}$). **b**, summary of Marchantia Mp*arf1* complementation assay using MpARF1 with AD deleted (MpARF1$^{AD}$). Lines indicate the number of independent transgenic lines checked. **c**, Raincloud plot of thallus area measurements (in halves) of 10-day-old wild-type and Mp*arf1* plants complemented with MpARF1 or MpARF1$^{AD}$ grown in mock (DMSO) or auxin (3 µM NAA). n=15-20. Statistical groups are determined by Tukey's Post-Hoc test ($p<0.05$) following one-way ANOVA. **d**, pictures of representative plants from **c**. Scale bar, 5 mm.

### Supplementary Figure 4

a

| Construct | Lines | Postembryonic rescue (%) | |
|---|---|---|---|
| | | Yes (full) | No |
| *pARF5:ARF5* | 5 | 100 | 0 |
| *pARF5:ARF5*$^{F333V,\ Y343S}$ | 5 | 100 | 0 |
| *pARF5:ARF5*$^{E372Q}$ | 4 | 100 | 0 |
| *pARF5:ARF5*$^{K378A}$ | 8 | 100 | 0 |
| *pARF5:ARF5*$^{K378M}$ | 7 | 100 | 0 |
| *pARF5:ARF5*$^{K378R}$ | 5 | 100 | 0 |

b

| Construct | Lines | Thallus growth rescue (%) | | Auxin responsivity rescue (%) | |
|---|---|---|---|---|---|
| | | Yes | No | Yes | No |
| *pMpARF1:MpARF1* | 6 | 100 | 0 | 100 | 0 |
| *pMpARF1:MpARF1*$^{F312V,\ Y322S}$ | 5 | 100 | 0 | 100 | 0 |
| *pMpARF1:MpARF1*$^{E351Q}$ | 3 | 100 | 0 | 100 | 0 |
| *pMpARF1:MpARF1*$^{R357A}$ | 4 | 100 | 0 | 100 | 0 |

c

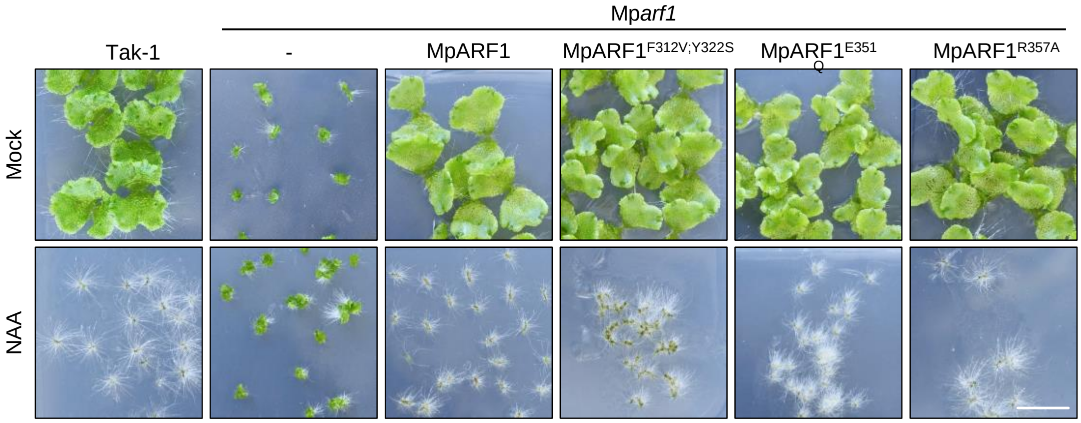


**Supplementary figure 4. AD putative hydrophobic cage perturbation does not affect ARF function**. **a**, summary of Arabidopsis *mp* complementation assay using ARF5 with hydrophobic cage residue mutations. **b**, summary of Marchantia Mp*arf1* complementation assay using MpARF1 with hydrophobic cage residue mutations. Lines indicate the number of independent transgenic lines checked. **c**, pictures of 14-day-old wild-type and Mp*arf1* plants complemented with MpARF1 HC-mutants grown in mock (DMSO) or auxin (3 µM NAA). Scale bar, 1 cm.

#### Supplementary Figure 5

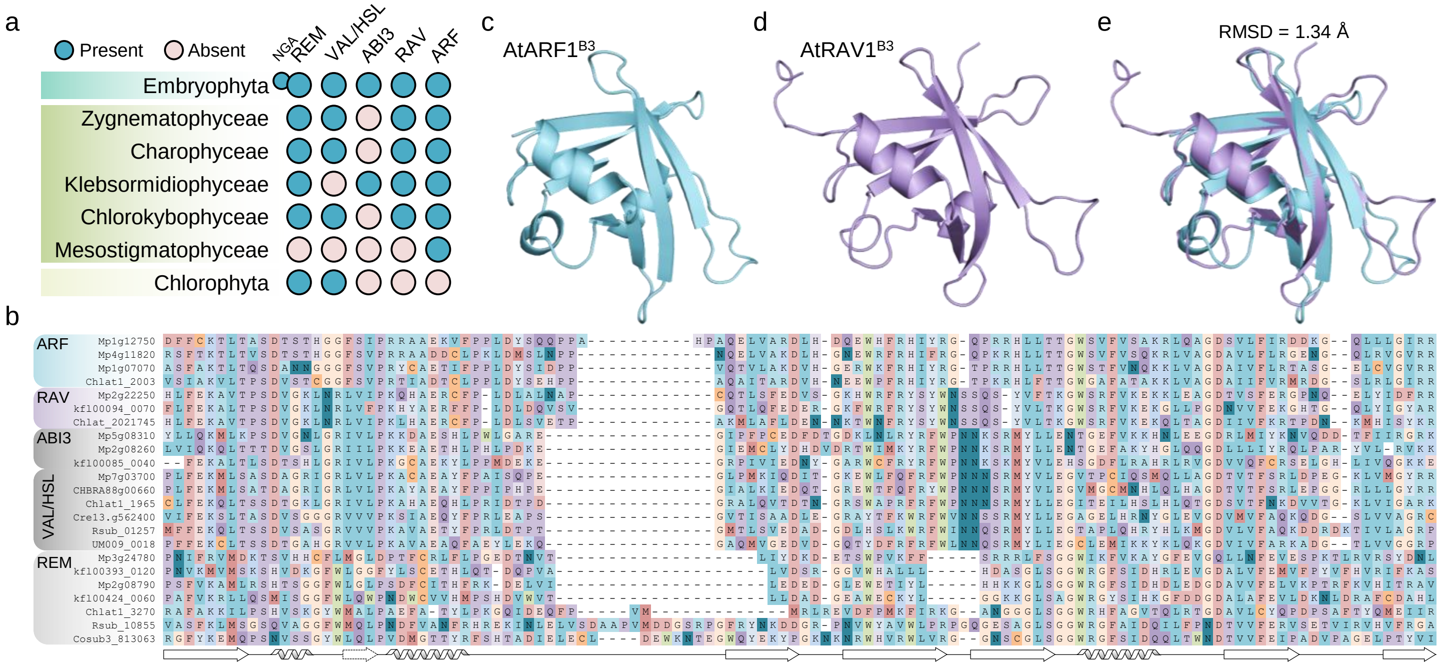


**Supplementary figure 5. ARF and RAV B3 homology**. **a**, Presence of B3 transcription factor subfamilies in the main Viridiplantae lineages as coloured circle representations. NGATHA (NGA) bubble represents a land plant specific expansion within the REM subfamily. **b**, Multiple sequence alignment of B3 domains from selected proteins. Different subfamilies are indicated on the left. Lower diagram is a schematic representation of the B3 secondary structure with seven β-sheets and two or three α-helixes. **c-d**, Structural comparison of AtARF1 and AtRAV1 B3s from crystal structures (PDB: 4ldx and 1wid in **c** and **d**, respectively) and their structure alignment in **d**.

**Supplementary Figure 6**

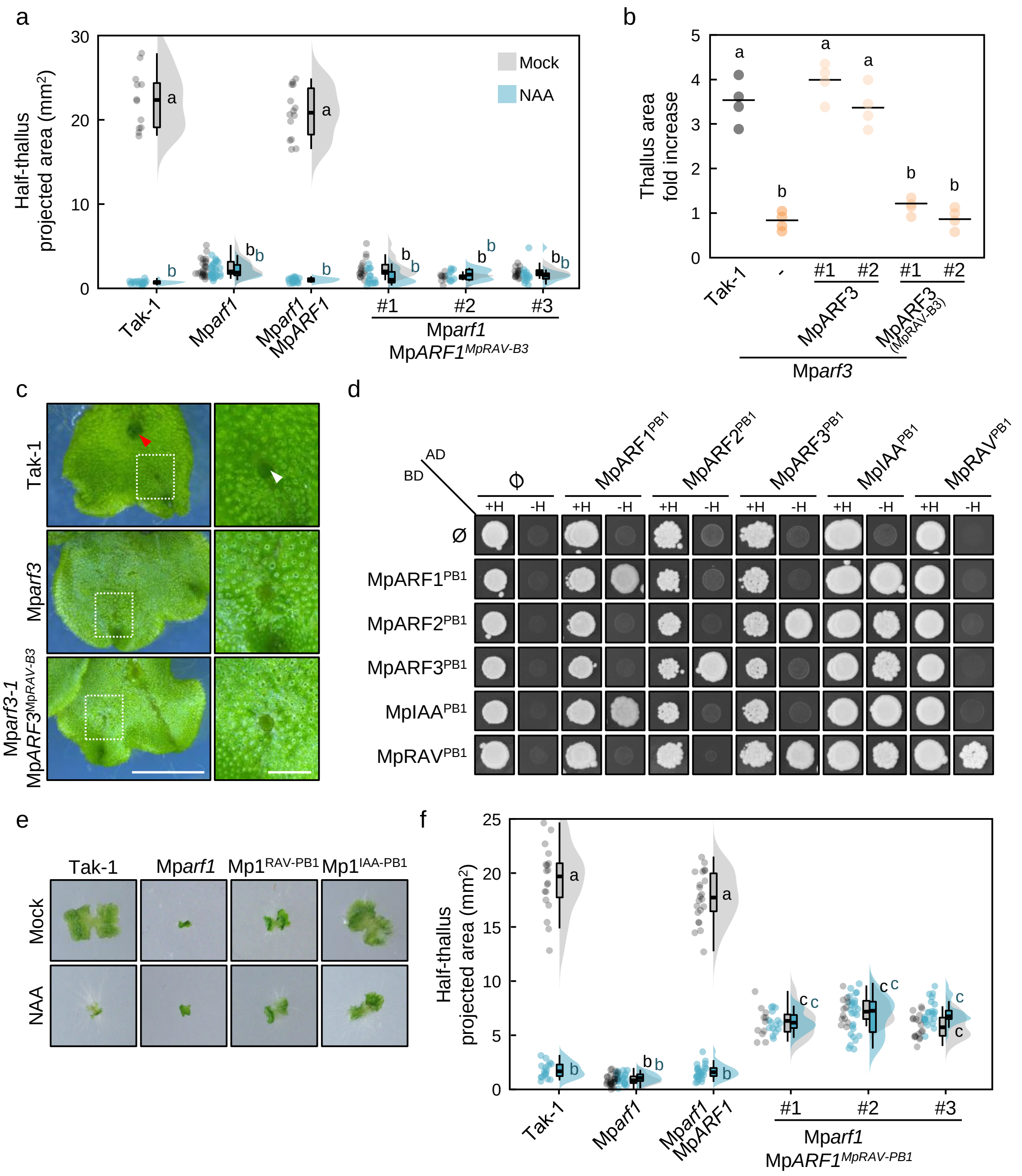

**Supplementary figure 6. RAV-ARF homologous domains are functionally divergent**. **a**, Raincloud plot of half-thallus area measurements of 10-day-old Mp*arf1* mutant complemented with different MpARF1 B3 domain swapped for MpRAV B3 lines grown in mock (DMSO) or auxin (3 µM NAA). Controls and line #1 data are reproduced from Fig. 2c. **b**, Dot plot of notch-driven thallus growth measured as projected area fold-change 14 days after excision (percentage of times day 0 area) in wild-type and Mp*arf3* plants. MpARF3 chimera bearing MpRAV B3 is expressed under the control of the endogenous MpARF3 promoter in the Mp*arf3* mutant. **c**, pictures of Tak-1, Mp*arf3* and Mp*arf3* complemented with a MpARF3 chimera carrying MpRAV B3 domain grown for one month after cutting. Left panels show apical region, and right panels inset of young gemma cups. Scale bar, 5 mm and 1 mm, respectively. Arrowhead points to developing gemma in the wild-type Tak-1 as opposed to empty cups associated with Mp*arf3* mutation. **d**, Yeast two-hybrid drop assay assessing pairwise Marchantia PB1 interactions. e, Pictures of 10-day-old plants shown in Fig. 2h. **f**, Raincloud plot of half-thallus area measurements of 10-day-old Mp*arf1* mutant complemented with additional MpARF1 PB1 domain swapped for MpRAV lines. Controls and line #1 data are reproduced from Fig. 2h.

### Supplementary Figure 7

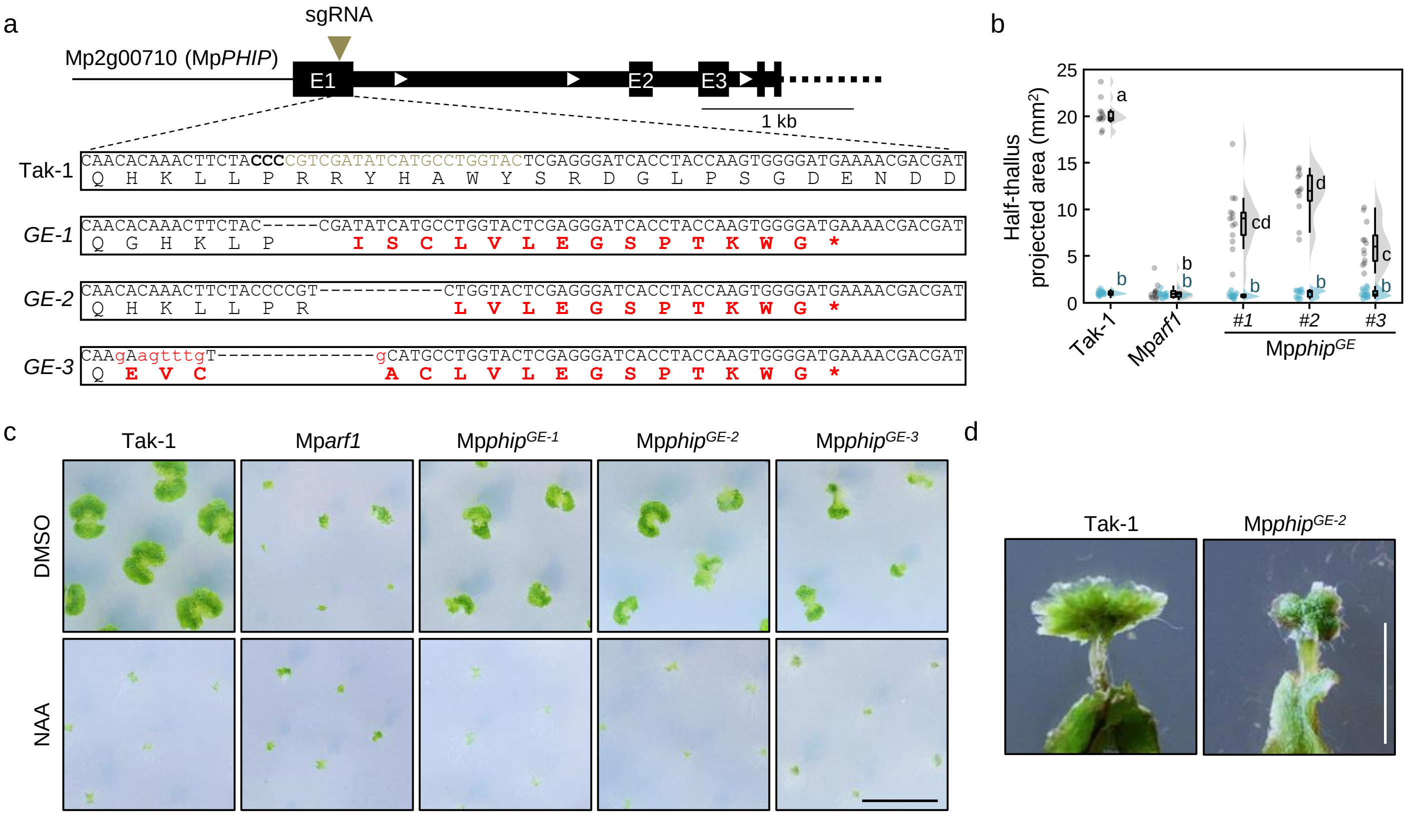


**Supplementary figure 7. MpPHIP genome editing design**. **a**, design of CRISPR/Cas9 mediated genome edited MpPHIP. Upper scheme represents the upstream region of Mp2g00710 loci with the gene CDS annotated (E, exon). Lower boxes show wild-type region targeted by Cas9 and the corresponding region in the genome edited lines aligned, with the ORF translated below nucleotide sequence. Red amino acids indicate different residues compared to wild-type ORF. **b**, Raincloud plot of thallus area measurements (in halves) of 10-day-old wild-type and *phip* mutant plants grown in mock (DMSO) or auxin (3 µM NAA). Controls and line GE-1 data are reproduced from Fig. 3b. **c**, Pictures of plants in **b**. Scale bar, 1 cm. **d**, Antheridophore pictures of 40-day-old plants after induction with far-red light for two weeks. Scale bar, 5 mm.

#### Supplementary Figure 8

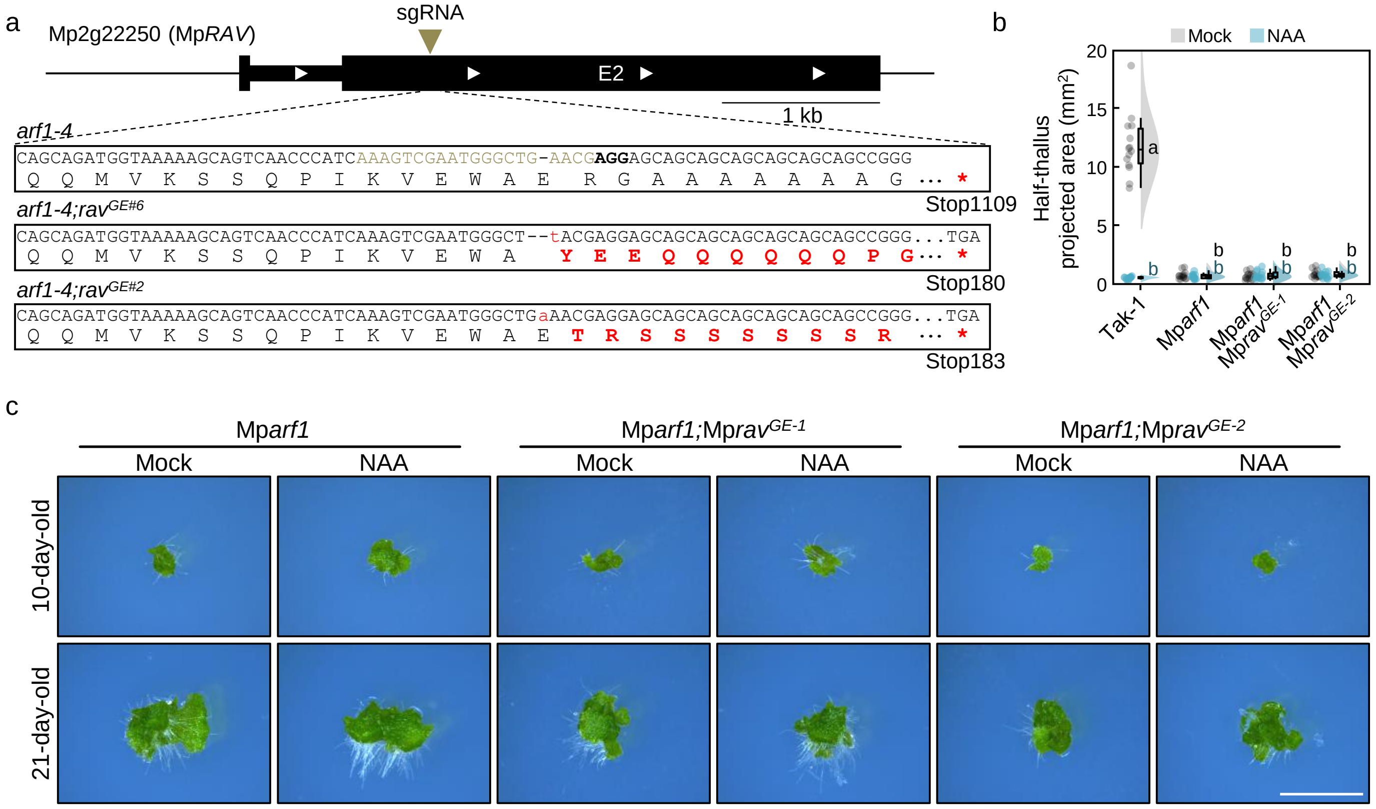


**Supplementary figure 8. MpRAV genome editing design**. **a**, design of CRISPR/Cas9 mediated genome editing of MpRAV. Upper scheme represents Mp2g22250 loci with the CDS annotated (E, exon). Lower boxes show wild-type region targeted by Cas9 and the corresponding region in the genome edited lines aligned, with the ORF translated below nucleotide sequence. Red amino acids indicate different residues compared to wild-type ORF. Stop indicates position of the first stop codon following the main ORF **b**, Raincloud plot of thallus area measurements (in halves) of 10-day-old wild-type and Mp*arf1;rav* mutant plants grown in mock (DMSO) or auxin (3 µM NAA). Controls and line GE-1 data are reproduced from Fig. 3f. **c**, Pictures of representative plants in **b**, upper panel, and same lines grown for 21 days, lower panels. Scale bar, 5 mm.

#### Supplementary Figure 9

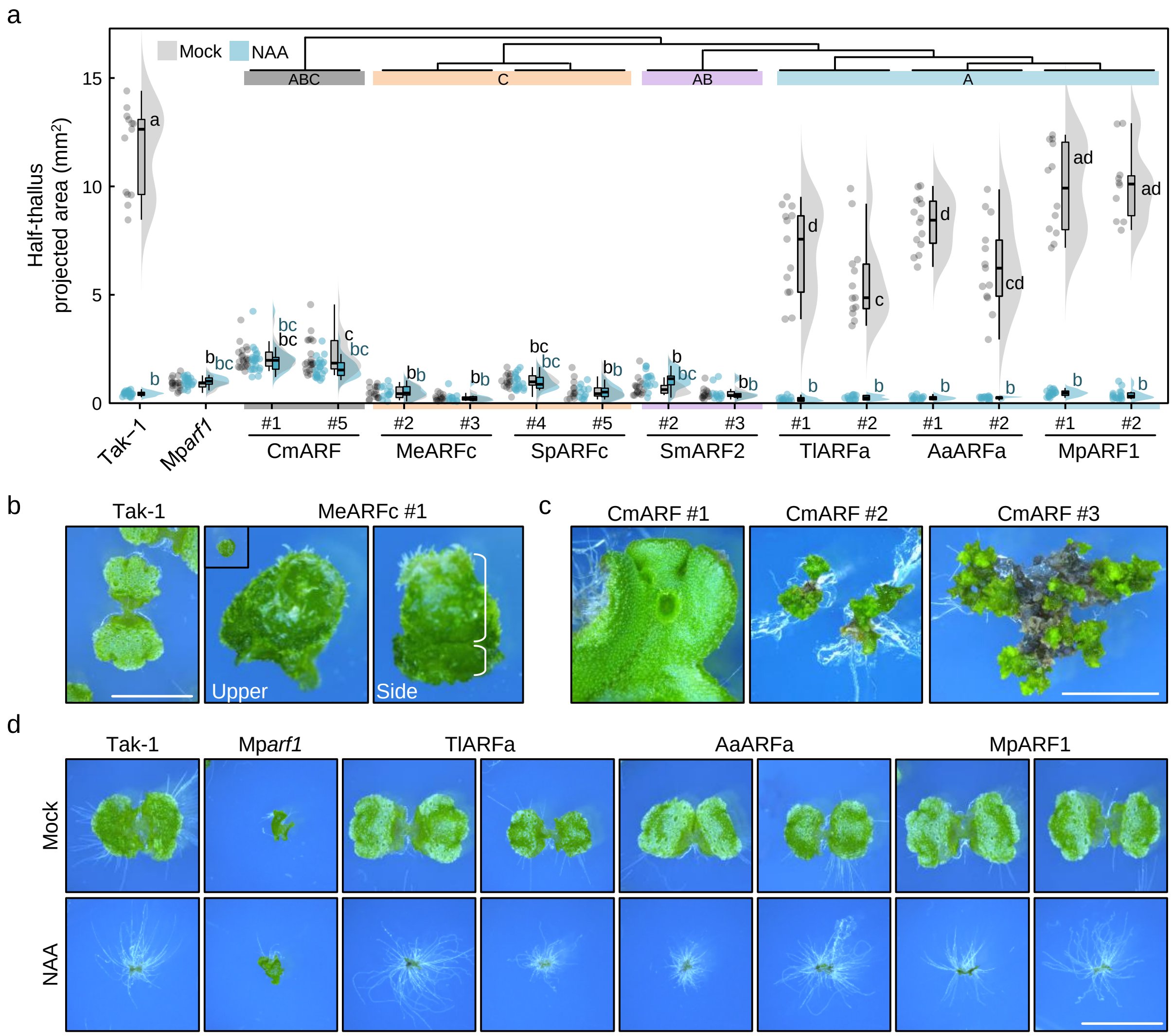


**Supplementary figure 9. Heterologous complementation of Mp*arf1* mutant**. **a**, Raincloud plot of thallus area measurements (in halves) of 10-day-old wild-type and mutant rescue lines grown in mock (DMSO) or auxin (3 μM NAA). **b**, pictures of a 10-day-old wild-type plant (left panel) compared to an aberrant-growing gemma forming a gemma cup (inset in middle panel is same scale as Tak-1) commonly found among Mp*arf1*;MeARFc lines. Upper and side views are shown. Right panel highlights the original gemma (lower part) and the cup formed from it. Scale bar, 5 mm. **c**, comparison of different Mp*arf1* lines expressing CmARF grown for 50 days after thallus transformation and selection. Lines 2 and 3 do not form clear tissues or gemma. Scale bar, 4 mm. **d**, Representative pictures of 10-day-old Mp*arf1* plants complemented with A-class ARFs from different bryophytes. Tl, *Takakia lepidozioides*; Aa, *Anthoceros agrestis*. Scale bar, 5 mm.

#### Supplementary Figure 10

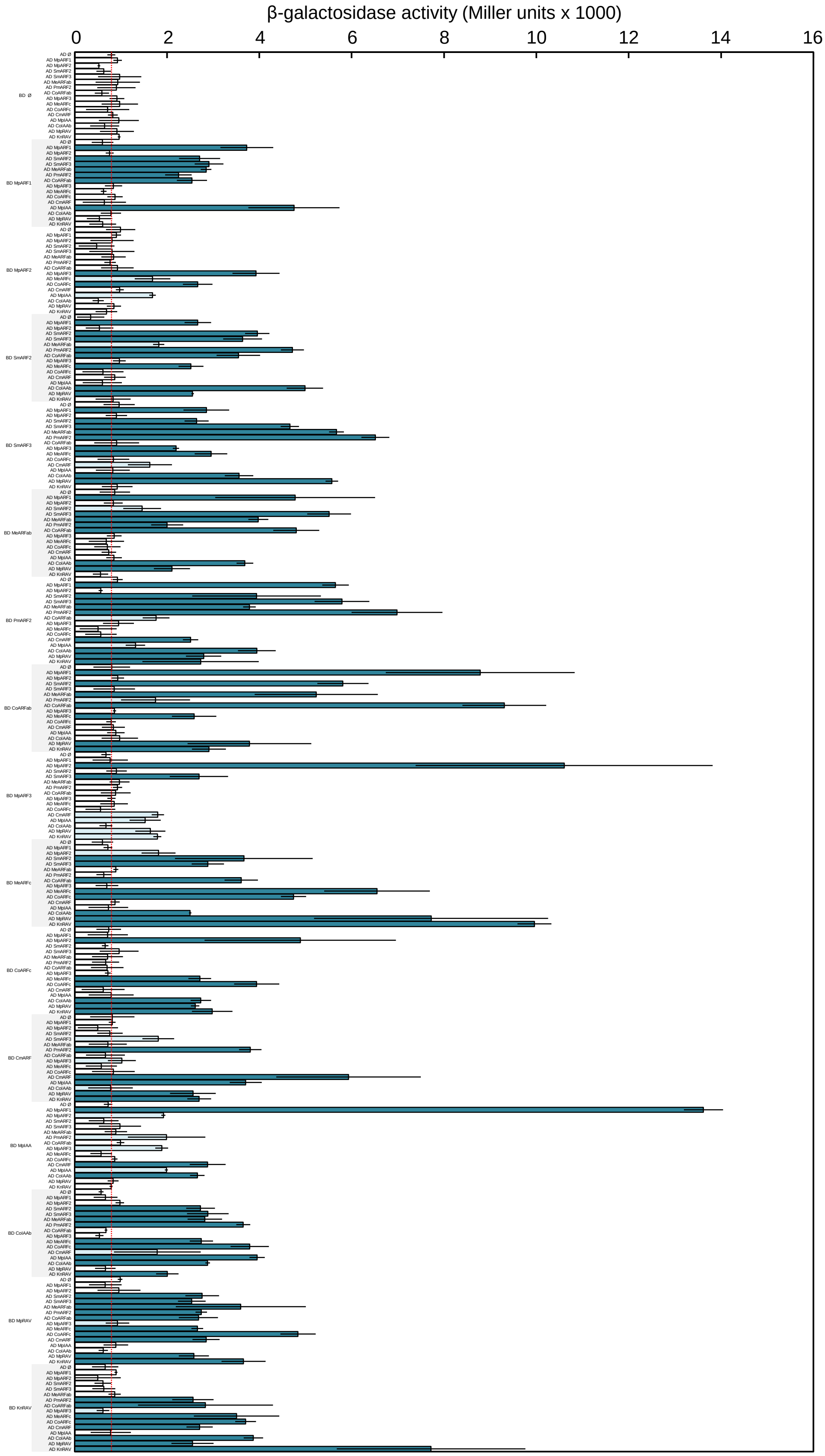


**Supplementary figure 10. Yeast two-hybrid pairwise PB1 interactions quantification by galactosidase assay**. Data represents absolute galactosidase quantified from two replicates of three independent lines. Bars indicate average of mean in replicates and error bars standard deviation. Red line represents control average value. Strong (dark cyan) and weak (light cyan) interactions are shown as average values with fold-change above one or two compared to control average.

#### Supplementary Figure 11

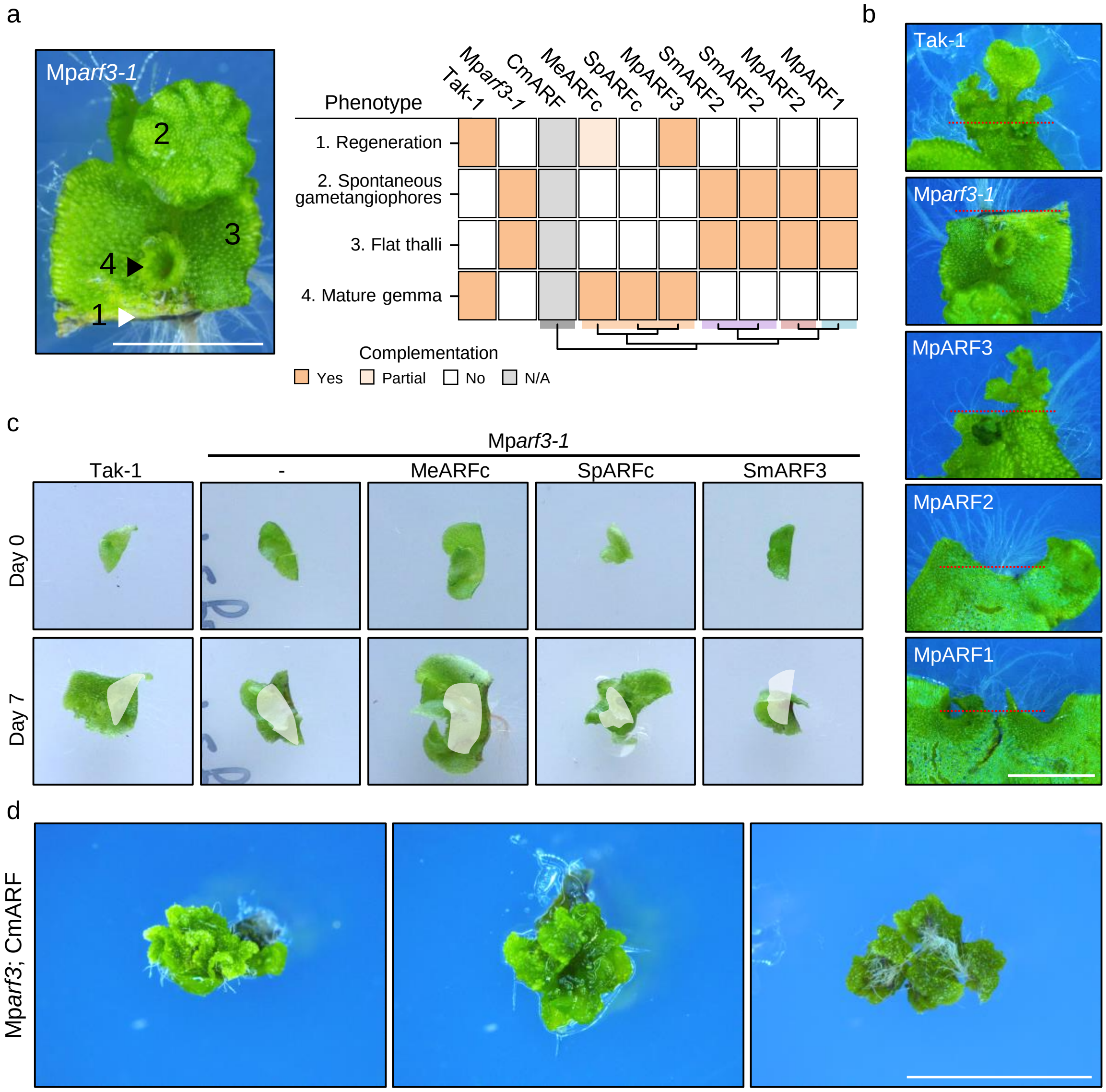


**Supplementary figure 11. Heterologous complementation of Mp*arf3* mutant**. **a**, Summary of Mp*arf3*-related phenotypes complementation using heterologous ARF sequences. Left panel depicts an apical sectioned region of an Mp*arf3* plant grown for 21 days showing the four accounted phenotypes. Black arrowhead points to the empty gemma cup, and white arrowhead indicates lack of thallus regeneration after excision. Right panel summarizes effect of ARF expression in the mutant. Partial complementation indicates instances of thallus regeneration but uncommon. N/A, for CmARF, indicates inability to determine (see **d**). **b**, Regeneration of Mp*arf3* mutants complemented with MpARFs 21 days after excision. **c**, Pictures of Mp*arf3* mutant complemented with algal ARFs. ARF coding sequences are expressed under the control of the endogenous MpARF3 promoter in the Mparf3-1 mutant. Upper row represents 20-day-old apical notches right after excision from adult plants. Lower row are the same plants after seven days of re-growth. White-shaded area in lower row is the same area occupied at day 0. **d**, Mp*arf3* lines expressing CmARF grown for 80 days after thallus transformation and selection. Cm, *Chlorokybus melkonianii*; Me, *Mesotaenium endlicherianum*; Sp, *Spirogyra pratensis*; Sm, *Spirogloea muscicola*. Scale bars, 5 mm (all panels).

### Supplementary Figure 12

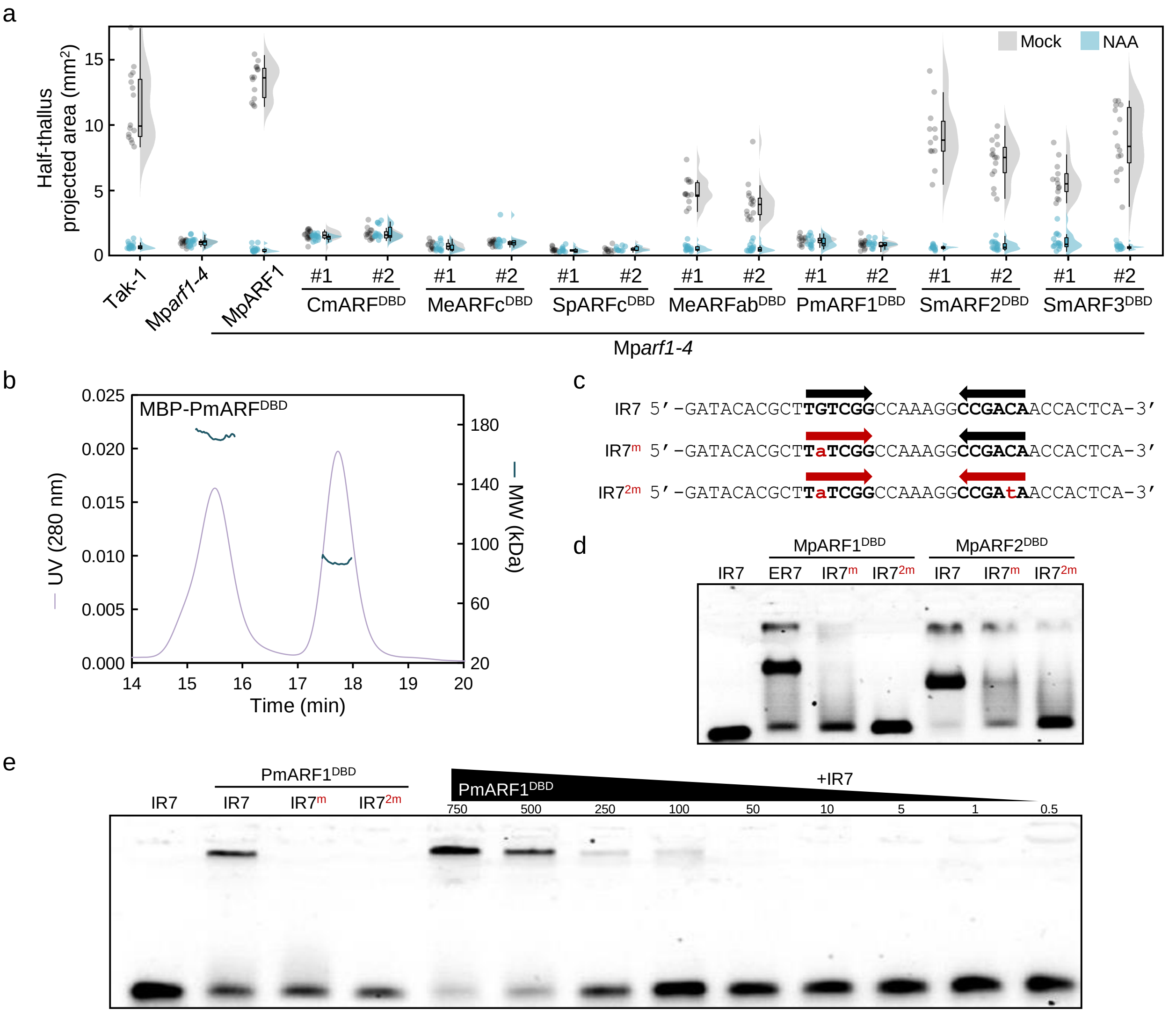


**Supplementary figure 12. AB-class ARF DNA binding conservation**. **a**, Raincloud plot of thallus area measurements (in halves) of 10-day-old wild-type and Mp*arf1* mutant complemented with MpARF1 lines grown in mock (DMSO) or auxin (3 μM NAA). n=15-25. **b**, Elution chromatogram showing PmARF$^{DBD}$ protein dimer formation in solution. Molecular weight calculated by size exclusion chromatography-multi-angle light scattering. Left and right peaks represent dimeric (172.3 kDa) and monomeric (87.5 kDa) forms, respectively. **c**, Probes used in EMSA assays. Only one strand is shown, with the AuxRE indicated. Red, lower case basepairs indicate mutated AuxRE. **d**, EMSA assay showing MpARF1 and MpARF2 DNA binding domains interaction with a bipartite ARF-specific binding site (IR7, inverted repeat), and the same elements mutated in one or two of the bipartite sites (IR7$^{m}$ and IR7$^{2m}$, respectively). **e**, EMSA assay showing PmARF1 DNA-binding domain interaction with the same binding site as in **d**. Right-most lanes show a PmARF1$^{DBD}$ titration experiment indicating concentration-dependent complex-formation. Numbers indicate PmARF1$^{DBD}$ concentration in nanomoles. Sm, *Spirogloea muscicola*; Pm, *Penium margaritaceum*.

#### Supplementary Figure 13

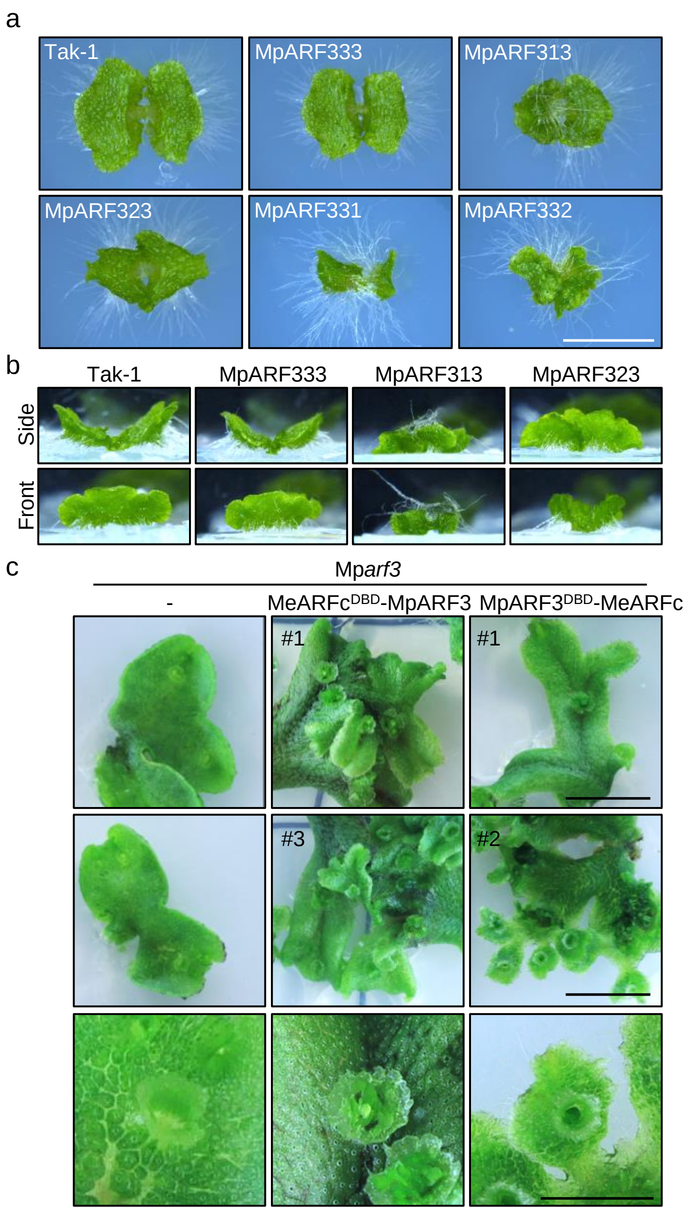


**Supplementary figure 13. Main C-class function conservation is encoded in the DNA binding domain**. **a**, Pictures of 10-day-old Mp*arf3* complemented lines using MpARF chimeras indicated in Fig. 7d. Scale bar, 5 mm. **b**, Detail of MpARF3 chimeras with MpARF1 and MpARF2 middle regions. Upper panels are side views and lower panels front views indicating differences with wild-type phenotypes. Plants and scale are the same as depicted in **a**. **c**, Pictures of Mp*arf3* mutant complemented with different MeARFc domains. First and second rows show plants grown from excised apical notches 30 after cutting. Scale bar, 1 cm. Third row are zoom-ins of gemma cup showing gemma formation in complemented lines. Scale bar, 2 mm. Me, *Mesotaenium endlicherianum*.
